# Dynamic Structure-based Pharmacophore Models for Virtual Screening of Small Molecule Libraries Targeting the YB-1

**DOI:** 10.1101/2023.07.26.550723

**Authors:** Lalehan Oktay, Ehsan Sayyah, Serdar Durdağı

## Abstract

In drug discovery, ligand-based techniques offer rapid screening, whereas structure-based approaches provide deeper insights but are time-consuming. Hybrid methods like structure-based pharmacophore models combine advantages for accurate screening of large ligand libraries. However, there are substantial limits to build structure-based pharmacophore models. Static models relying on a single co-crystallized structure or docking pose often fall short in capturing the dynamic nature of binding interactions. In this study, we present dynamic structure-based pharmacophore models, aimed at better representing physiological conditions and addressing these challenges. The urgent need for improved cancer treatment has led to the search for new chemotherapeutic strategies. Y box binding protein 1 (YB-1) is a multifunctional protein associated with tumor progression and treatment resistance in various cancers. For the first time in the literature, our study utilizes a known small molecule YB-1 inhibitor (SU056) bound to the active regions of the RNA-binding sites to develop dynamic structure-based pharmacophores. These models were then used in the screening of large ligand libraries.

## 1. Introduction

Ligand-based techniques, such as 3D pharmacophore modeling, are useful for rapidly screening small molecule libraries in large databases due to their speed. The Pharmacophore model is represented as a set of pharmacophore points, each characterized by the following features: type of functional group, point center, and spread (α).^1^ Each pharmacophore point is modeled as a three-dimensional (3D) spherical Gaussian volume represented by Cartesian coordinates and spread. The definition of a Gaussian volume is as follows:

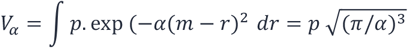

In this equation, *V*_*α*_ represents the atomic Gaussian volume; *p* is the normalization constant used to relate the total volume to atomic volumes; *m* denotes the center of the Gaussian; and *r* is the integrated distance variable. 3D structures can be utilized to identify the molecular properties required for a protein to bind to a ligand. These fingerprints can be labeled through the calculation of physicochemical parameters, including hydrophobicity (logP), topological polar surface area, hydrogen bond acceptor (A) and donor (D) groups, and ring-based parameters (R).

On the other hand, structure-based approaches can provide deeper insights into the structure of the target protein’s binding site, making them beneficial for identifying a broader range of active compounds, but they are generally more time-consuming compared to ligand-based methods. Hybrid methods, like structure-based pharmacophore models, can combine the advantages of both ligand and target-based approaches by creating energy-optimized, structure-based pharmacophores, enabling fast and accurate screening of ultra-large ligand libraries.^2^ Nonetheless, there are notable constraints when it comes to constructing structure-based pharmacophore models. Firstly, despite the Protein Data Bank (PDB) containing over 207,000 structures, only a small portion of them is in a ligand-bound holo form, with the majority being in an apo form. This disparity limits the availability of ligand-bound structures for generating comprehensive structure-based pharmacophore models. Secondly, even when a ligand is present in the binding site, there might be a scarcity of diverse ligands accessible for the target protein, leading to restricted and unvarying possibilities for formulating structure-based pharmacophore hypotheses. Lastly, it is evident that static structure-based pharmacophore models, generated solely based on one co-crystallized structure or docking pose, are likely to yield limited success.

Hence, there arises a necessity for dynamic structure-based pharmacophore models that can better capture the complexities of physiological conditions. To address this, in the current study, extensive long (1 μs) all-atom molecular dynamics (MD) simulations are carried out on a model protein-ligand system (Y-box binding protein 1 (YB1)/inhibitor). Throughout these simulations, numerous trajectories are recorded (e.g., 1000 trajectories for each 1 μs), and structure-based pharmacophore models also known as E-pharmacophores are constructed for each snapshot, representing the protein-ligand complex at various time points.

Within the complex realm of cellular biology, various proteins play vital roles in regulating gene expression and maintaining cellular homeostasis. Among these, the YB-1 stands out as a multifunctional, evolutionarily conserved nucleic-acid-binding protein with a cold-shock domain (CSD).^3^ YB-1 has been implicated in transcriptional and translational regulation, drug resistance, and cellular stress responses to extracellular signals.^3^ Mainly, the highly conserved YB-1 CSD domain detects both RNA and DNA.^4^ Notably, CSD-containing proteins are active in various RNA metabolism processes due to their ability to bind both DNA and RNA.^5^ The YB-1 CSD is made up of five-antiparallel β-barrels and capacitates YB-1 to bind to single-stranded DNA and RNA, particularly at lower temperatures. It comprises two RNA-binding motifs known as RNP-1 and RNP-2 which are on the β2 and β3, respectively.^6^ The CSD is structurally like the major cold shock proteins found in bacteria.^7,8^ In bacteria, cold shock proteins play essential roles in cold adaptation by acting as RNA chaperones.^9^ They are involved in the regulation of gene expression, including the control of the cold-shock stress response and the translational masking of messenger RNA.^4^ The YB-1 cold shock domain, in particular, has been structurally characterized and shown to have a high degree of similarity to bacterial cold shock proteins.^10^ Overall, the cold shock domain is a versatile and evolutionarily ancient domain that plays important roles in nucleic acid binding and regulation of gene expression in response to cold stress. NMR solution studies revealed that residues Trp15, Phe24, Phe35, and His37 on the exterior of the barrels are part of the nucleic-acid-binding domain.^6^

YB-1 is primarily localized in the cytoplasm but can translocate to the nucleus in response to certain stimuli.^11^ In the cytoplasm, YB-1 can bind to mRNAs or Y-box elements in the promoter regions of genes, regulating their translation and transcription.^12^ In the nucleus, YB-1 acts as a transcription and splicing factor, modulating the expression of its target genes.^13^ YB-1 has been implicated in various physiological and pathological processes, including cell proliferation, fatty acid synthesis, antifibrotic effects, and cancer progression.^3,13-15^ Additionally, YB-1 has been found to interact with other proteins, such as Elongin BC and transportin 1, which can affect its stability, assembly of protein complexes, and subcellular localization.^16,17^ Overall, YB-1 is a multifunctional protein that plays a critical role in various cellular processes and has implications in disease development and progression.

In a recent study identifying a potent YB-1 inhibitor, Tailor *et al*.^18^ discovered an azopodophyllotoxin small molecule called SU056, which effectively inhibits tumor growth and progression by targeting YB-1.^18^ The YB-1 inhibitor demonstrated various beneficial effects, including inhibiting cell proliferation, reducing resistance to apoptosis in ovarian cancer cells, and causing cell cycle arrest in the G1 phase.^18^ Treatment with SU056 led to changes in protein expression associated with apoptosis and RNA degradation pathways, while downregulating the spliceosome pathway. For the identification of the target protein for SU056, the Cellular Thermal Shift Assay (CETSA) is employed.^18^ CETSA is a method that uses the thermal stability of proteins to monitor the effect of ligand binding in live cells. Using CETSA, cells or tissue are subjected to 10 different temperatures all of which cause denaturation of proteins. When a ligand which enhances the thermal stability of a protein is bound, the protein will escape denaturation. Concentrations of the protein before and after CETSA are used to determine successful ligand binding. In their study, among the 804 soluble proteins, YB-1, TMSB10, SUMO2, PSMB2, TMSB4X, and CALM3 showed increased thermal stability.^18^ Further western blot analysis reveals that SU056 actively inhibits YB-1, in particular.^18^ Most importantly, SU056 sensitized ovarian cancer cells to paclitaxel.^16^ Dose-dependent MTT assays show the synergistic effect of combinatorial treatment with SU056 and paclitaxel. Furthermore, taxol efflux was also significantly reduced with SU056-added treatment than in paclitaxel only treated cells.^18^ Overall, their results suggest that inhibiting YB-1 could be a promising strategy to reduce ovarian cancer progression, counteract treatment resistance, and potentially improve patient outcomes.^18^

## 2. Methods

### 2.1 Molecular Docking and Molecular Dynamics (MD) Simulations of YB-1/SU056 complex

The structure of YB1 with the PDB ID 6KUG was obtained and prepared using the Protein Preparation tool within the Maestro molecular modeling suite.^19^ The preparation process included adding hydrogens and creating disulfide bonds. PROPKA was employed to determine the protonation states of the residues at neutral pH.^20^ Subsequently, restrained minimization was performed using the OPLS3 force field, with a convergence criterion set at a 0.3 Å cutoff value for the RMSD of heavy atoms.^21^

The center of masses of previously reported RNA-binding residues Phe85-Lys118, His87-Glu121, Trp65-Phe74-Asp83 and Trp65-Asn67 were used as the four grid box centers.^22^ Molecular docking of YB-1 (PDB ID, 6KUG)^22^ to YB-1 inhibitor SU056^16^ was conducted using the Glide/SP docking algorithm at the four grid boxes. SU056 was prepared before the docking using LigPrep module of the Maestro molecular modeling tool. OPLS3 forcefield with neutral pH was used in ligand preparation.^21^ Conducting all-atom MD simulations for the YB-1/SU056 complexes at the four grid coordinates involved running simulations of 1 μs duration each using Desmond. The top-docking poses were used to initialize the simulations. Nose-Hover and Martyna-Tobias-Klein approaches were used at 310 K and 1.01325 bar for the thermostat and barostat throughout the simulations.^23,24^ To generate the complex structures for the simulations, they were immersed in a water box containing counter ions and TIP3P waters, extending 10 Å from the protein’s edges. A default protocol was followed for the geometry optimization and equilibrium. Each simulation produced 1000 trajectory frames throughout the 1 μs production run, which were subsequently employed in the generation of structure-based pharmacophores.

### 2.2 Construction of Dynamic Structure-based Pharmacophore Modeling

Dynamic structure-based pharmacophore hypotheses were derived from the 1 μs all-atom MD simulations of YB-1 and SU056 complexes bound to four different conserved RNA-binding sites: Phe85-Lys118, His87-Glu121, Trp65-Phe74-Asp83, and Trp65-Asn67. For each of the 1000 frames extracted from the MD trajectories, pharmacophore hypotheses with 3, 4, and 5-features were generated using the e-Pharmacophore module of Phase within the Maestro molecular modeling package. The e-Pharmacophore approach involves mapping the rescored Glide/XP (extra precision) energetic descriptors of the receptor-ligand complex onto the pharmacophore sites, allowing for quantification and ranking of the hypotheses.^2^ The most observed receptor-ligand-derived pharmacophore hypothesis in each of the trajectories is determined, and the frames exhibiting these pharmacophore features are ranked based on RMSD values to the average structure of the respective MD simulations. These selected optimized representative structures are used in grid generation and virtual screening.

### 2.3 Virtual Screening of Small-Molecule Libraries using Dynamic Pharmacophore Models

The LigPrep module of the Maestro was used in the preparation of the ligand files from small molecule libraries: (i) 300K Representative Compounds Library from ChemDiv (∼300.000 compounds) (https://www.chemdiv.com), (ii) Hit Locator Library from Enamine (∼460.000 compounds) (https://enamine.net), (iii) HTS Compound Collection from Life Chemicals (∼525.000 compounds) (https://lifechemicals.com) and (iv) Specs Screening Compound Library (∼350.000 compounds) (https://www.specs.net). Prepared ligands were screened with the most common e-Pharmacophore hypotheses using Phase. Molecules are ranked based on their alignment to the spatial arrangements of the features based on the RMSD of the sites, volume, vector alignments, which make up the Fitness score. The fitness score, which varies between -1.0 and 3.0, is determined by a linear combination of site and vector alignment scores, along with the volume score. The reference ligand, which represents an exact match with the hypothesis, achieves a flawless fitness score of 3.0.^1,2^ Molecules having Phase Fitness score over 2 are docked with the Glide/SP algorithm to the optimized protein structure from the chosen representative frame. Top-scoring protein-ligand complexes are subjected to 100 ns MD simulations and end-point MM/GBSA calculations.

## 3. Results

Generating e-Pharmacophore models that adapt to the dynamic binding pocket of the protein structure offers a comprehensive and refined understanding of the nature of the binding pocket. 1000 frames from 1 μs all atom simulations of YB-1/SU056 complexes at all of the four RNA-binding sites were retrieved and pharmacophore hypotheses with 3, 4 and 5 features were generated using the Phase e-Pharmacophore pharmacophore model development pipeline. The most observed 4-sited pharmacophore hypotheses among the 1000 trajectory frames of 1 μs MD simulations of the YB-1/SU056 complex for each of the four reported RNA-binding sites are given in Table 1 and Figure 1. Since the CSD RNA-binding site is superficial and shallow, we chose to proceed with 4-sited e-Pharmacophore hypotheses to define the binding site. Figure 2 depicts representation of dynamic pharmacophore model development for one of the grid sites. ADRR, AHRR, AHRR, and AHRR pharmacophore models were observed frequently throughout the simulations at Phe85-Lys118, His87-Glu121, Trp65-Phe74-Asp83, and Trp65-Asn67 sites, respectively.

**Table 1.** Dynamic structure-based pharmacophore hypotheses for each of the YB-1 CSD RNA-binding residues were searched. 1 μs MD simulations with SU056 for each reported 4 mRNA binding sites lead to discovery of most common pharmacophore hypothesis among 1000 frames of simulations. (top) Representative features (i.e., the frame that shows lowest RMSD to the average protein-ligand complex structure throughout the simulations) were considered. Top-panel at the table shows the frame number that represent lowest RMSD to the average complex structure. (bottom) Four libraries, were screened using the respective pharmacophore models at each of the four sites. Compounds that fit the pharmacophore hypothesis with “Fitness” score over 2 were used for docking to the newly generated grids from frames that showed above pharmacophore hypothesis and were closest to average frame in RMSD. Pharmacophore features, A: hydrogen-bond acceptor, D: hydrogen-bond donor, R: aromatic ring; H: hydrophobic.

| Phe85-Lys118 | His87-Glu121 | Trp65-Phe74-Asp83 | Trp65-Asn67 |
| --- | --- | --- | --- |
| ADRR; Frame No:690<sup>th</sup> | AHRR; Frame: 316<sup>th</sup> | AHRR; Frame: 808<sup>th</sup> | AHRR; Frame 630<sup>th</sup> |

| Libraries | Total compound count | Phe85-Lys118 | His87-Glu121 | Trp65-Phe74-Asp83 | Trp65-Asn67 |
| --- | --- | --- | --- | --- | --- |
|  |  | ADRR | AHRR | AHRR | AHRR |
| ChemDiv Representative BM Set | 300000 | 685 | 494 | 19 | 760 |
| Enamine HLL | 460000 | 1075 | 342 | 23 | 928 |
| Life Chemicals HTS | 525000 | 1459 | 413 | 15 | 1745 |
| SPECS SC | 350000 | 423 | 164 | 9 | 446 |

**Figure 1.**
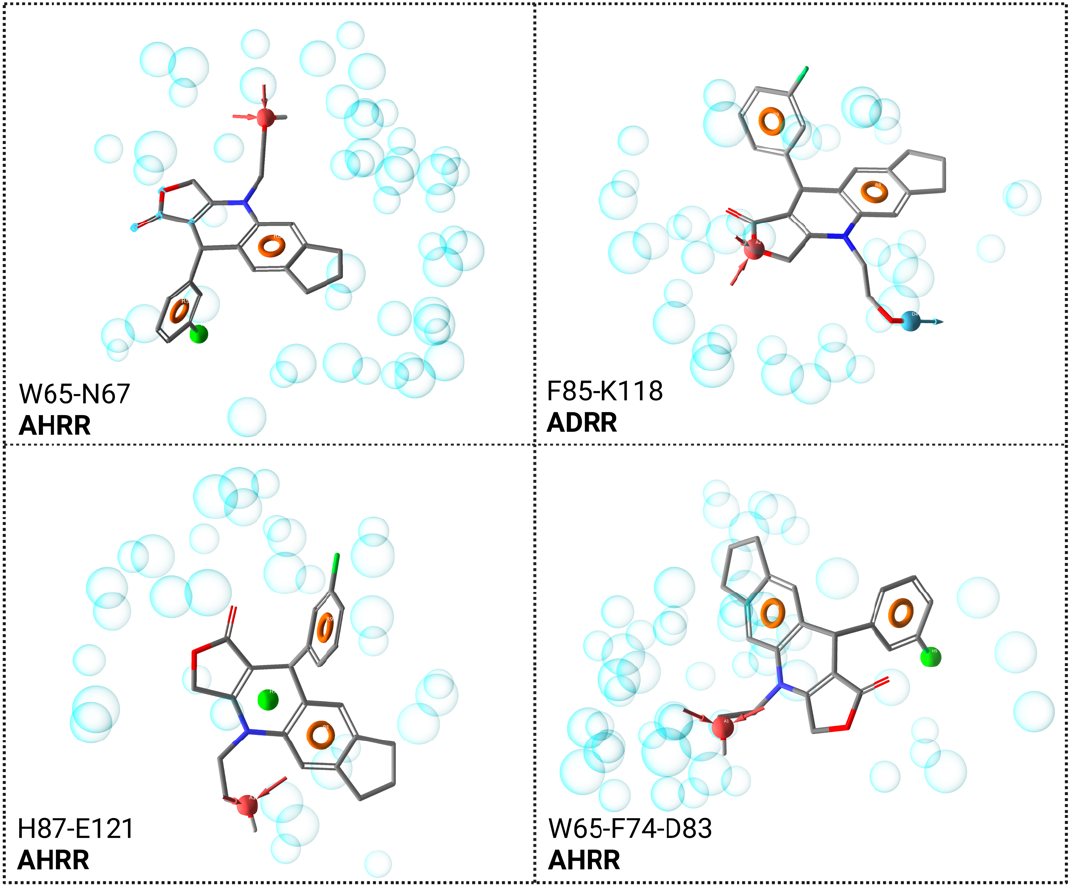
Dynamic structure-based e-Pharmacophore hypotheses of all 4 RNA-binding sites.

**Figure 2.**
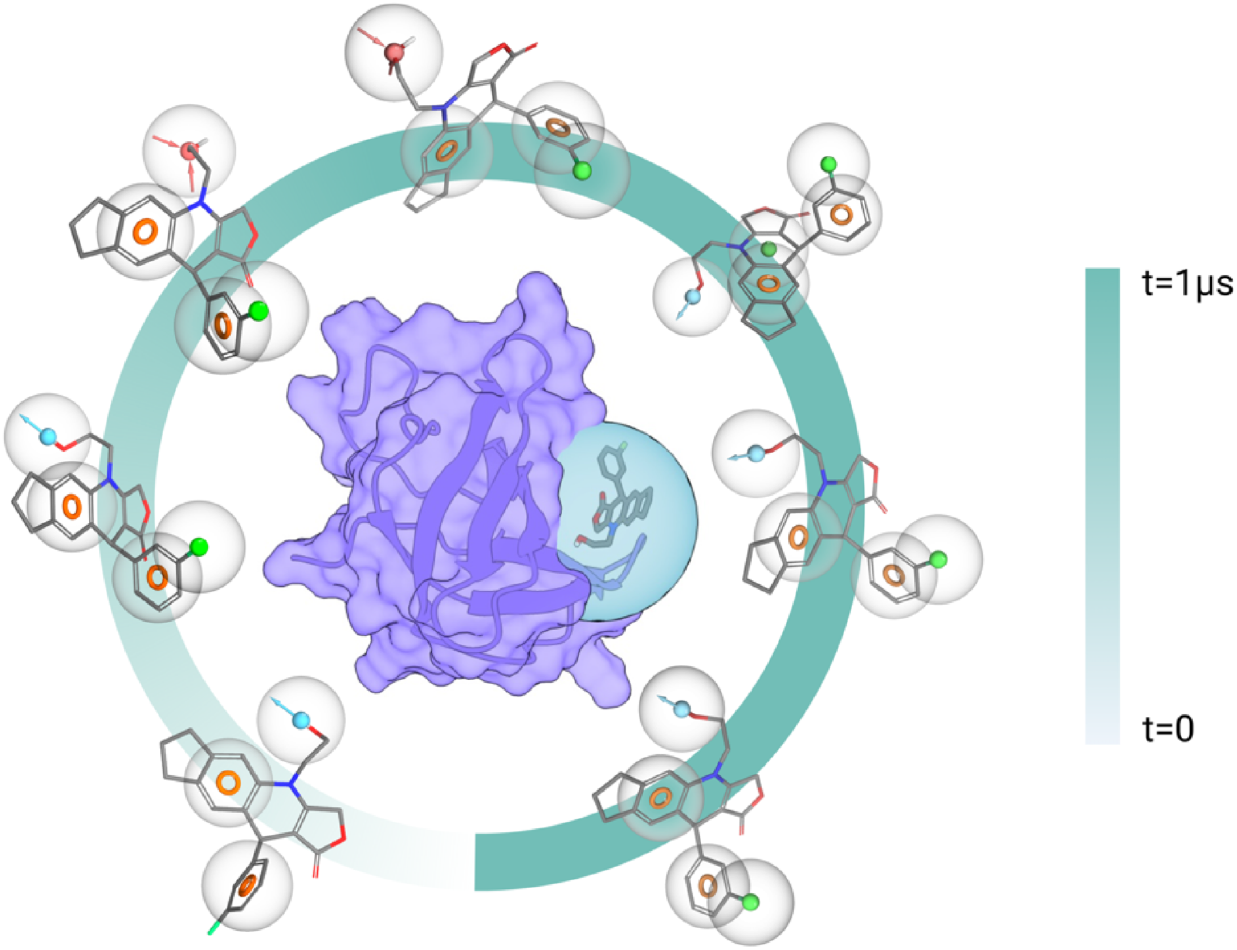
A known YB-1 inhibitor SU-056 was docked to the known crucial binding pockets of the target protein and 1 us MD simulations were conducted. Throughout these simulations, 1000 snapshots representing protein/ligand complexes were captured, and these poses were utilized in the development of e-Pharmacophores.

Using the optimized conformations of the protein structure and its respective ligand-receptor-derived pharmacophore hypothesis, screening of the selected 4 compound libraries: (i) 300k Representative Compounds Library from ChemDiv (∼300.000 compounds) (https://www.chemdiv.com), (ii) Hit Locator Library from Enamine (∼460.000 compounds) (https://enamine.net), (iii) HTS Compound Collection from Life Chemicals (∼525.000 compounds) (https://lifechemicals.com) and (iv) Specs Screening Compound Library (∼350.000) compounds) (https://www.specs.net)) were conducted. Only compounds that fit 3 out of 4 of the pharmacophore sites and had Phase Fitness score of over 2 were used for docking to the optimized YB-1 representative structure. (see Table 1 and Figure S1)

Frames exhibiting the chosen e-Pharmacophore hypotheses were selected and new grid centers were generated using the conformation of the protein structure at the specified trajectory frame. Selected compounds from each library were docked to the newly generated receptor grids. 100 ns all-atom MD simulations for the protein-ligand complexes and average MM/GBSA calculations over the trajectories were conducted. Results highlight expected interactions with the conserved RNA-binding sites. Table 2 represents 2D structures, library IDs, docking scores, fitness scores, and average MM/GBSA scores of selected top-scored hit compounds from each library. Ligand interaction diagrams of the top scoring compounds from each library and each binding site are summarized in Figure S2. Ligands bound to the conserved residues mostly comprise of fused ring systems and charged groups. The nature of the YB-1/ligand interactions mostly comprise of π-π stacking or π-cation interactions, mimicking the protein-RNA interactions as the CSD. Figure 3 illustrates the representative ligand interaction diagram of one of the chosen hits (i.e., AP-501/41001654 from Specs SC library) at the W65-N67 site. The compound demonstrates significant interactions, including π-π stacking with Phe74 and Phe85. Additionally, it forms crucial nonbonding interactions with Lys118, Gly118, and Glu121 (hydrogen bonding interactions with these residues), as well as hydrophobic contacts with Trp65 and Tyr72. The presence of these well-known key amino acid interactions in the chosen hit compounds strongly supports their potential inhibitory activity. In Figure S2, binding pocket residue-hit ligand interactions for all of the selected hits were depicted.

**Table 2.**
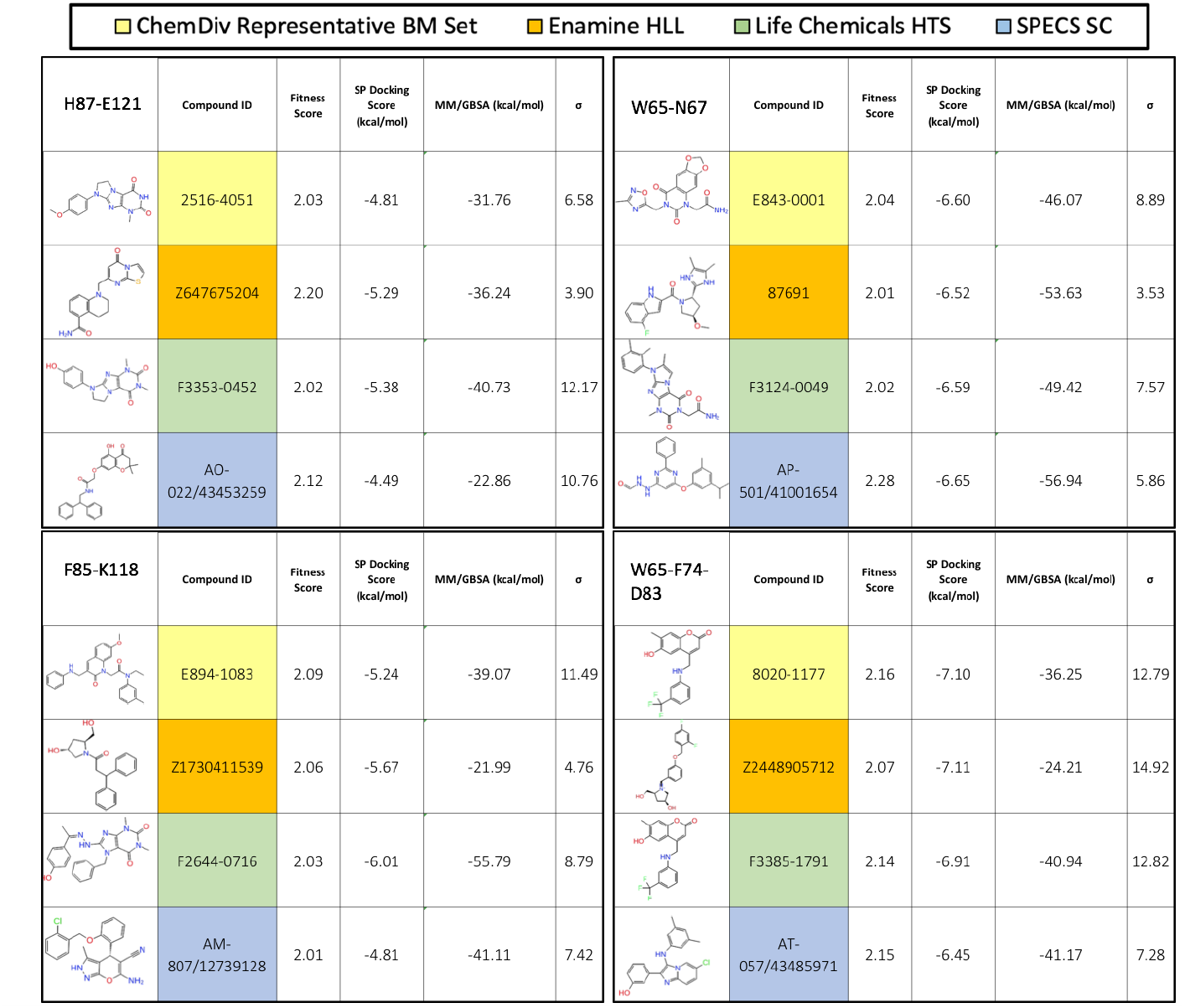
2D structures, library IDs, docking scores, fitness scores, and average MM/GBSA scores of selected top-scored hit compound from each library.

| ChemDiv Representative BM Set Enamine HLL Life Chemicals HTS SPECS SC |  |  |  |  |  |  |  |  |  |  |  |
| --- | --- | --- | --- | --- | --- | --- | --- | --- | --- | --- | --- |
| H87-E121 | Compound ID | Fitness Score | SP Docking Score (kcal/mol) | MM/GBSA (kcal/mol) | $\sigma$ | W65-N67 | Compound ID | Fitness Score | SP Docking Score (kcal/mol) | MM/GBSA (kcal/mol) | $\sigma$ |
|  | 2516-4051 | 2.03 | -4.81 | -31.76 | 6.58 |  | E843-0001 | 2.04 | -6.60 | -46.07 | 8.89 |
|  | Z647675204 | 2.20 | -5.29 | -36.24 | 3.90 |  | 87691 | 2.01 | -6.52 | -53.63 | 3.53 |
|  | F3353-0452 | 2.02 | -5.38 | -40.73 | 12.17 |  | F3124-0049 | 2.02 | -6.59 | -49.42 | 7.57 |
|  | AO-022/43453259 | 2.12 | -4.49 | -22.86 | 10.76 |  | AP-501/41001654 | 2.28 | -6.65 | -56.94 | 5.86 |
| F85-K118 | Compound ID | Fitness Score | SP Docking Score (kcal/mol) | MM/GBSA (kcal/mol) | $\sigma$ | W65-F74-D83 | Compound ID | Fitness Score | SP Docking Score (kcal/mol) | MM/GBSA (kcal/mol) | $\sigma$ |
|  | E894-1083 | 2.09 | -5.24 | -39.07 | 11.49 |  | 8020-1177 | 2.16 | -7.10 | -36.25 | 12.79 |
|  | Z1730411539 | 2.06 | -5.67 | -21.99 | 4.76 |  | Z2448905712 | 2.07 | -7.11 | -24.21 | 14.92 |
|  | F2644-0716 | 2.03 | -6.01 | -55.79 | 8.79 |  | F3385-1791 | 2.14 | -6.91 | -40.94 | 12.82 |
|  | AM-807/12739128 | 2.01 | -4.81 | -41.11 | 7.42 |  | AT-057/43485971 | 2.15 | -6.45 | -41.17 | 7.28 |

**Figure 3.**
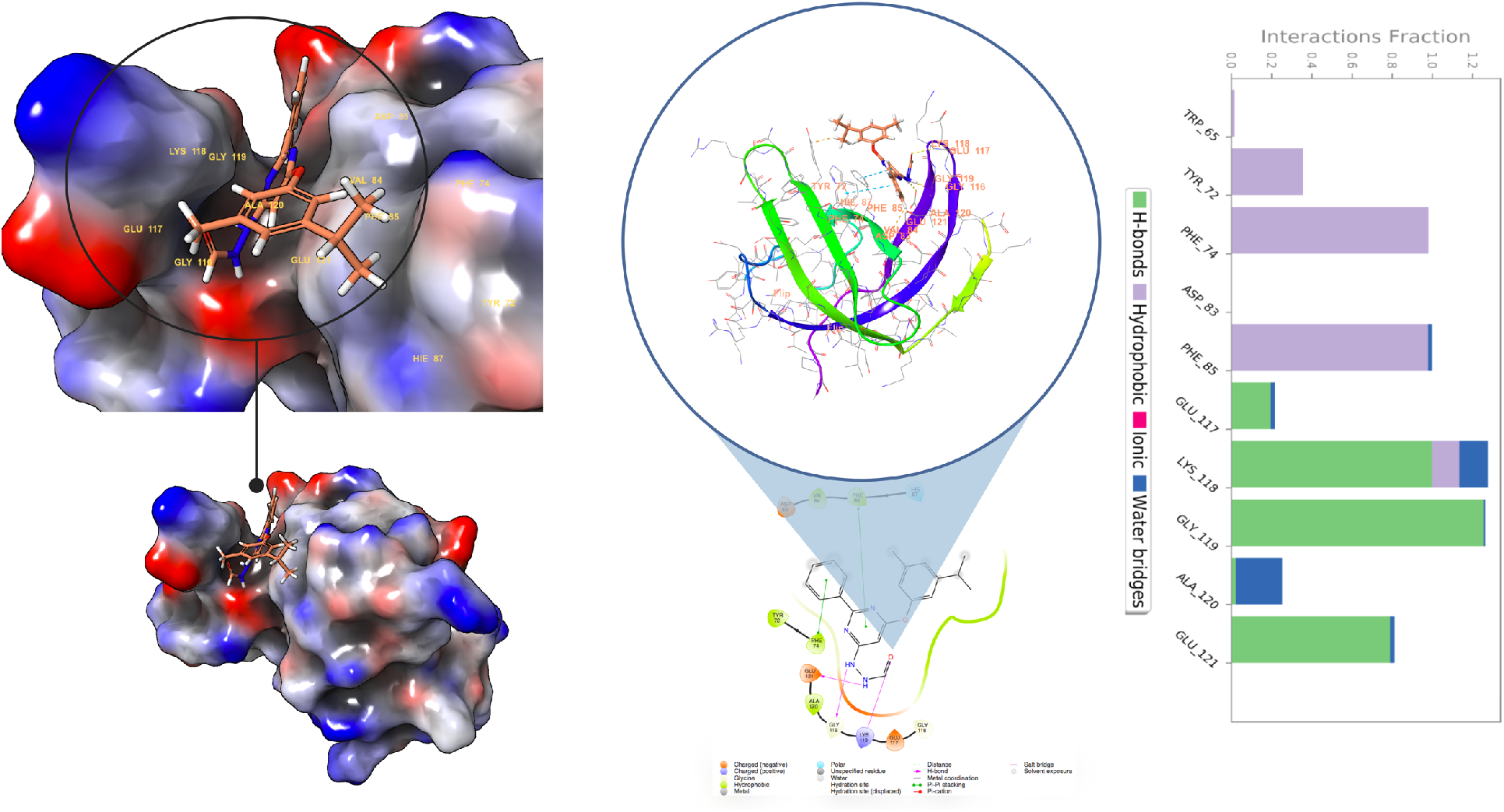
3D and 2D ligand interactions diagrams of selected hit compound (AP-501/41001654) as a proposed new YB1 inhibitor from Specs SC library at the W65-N67 site. Figure also shows interaction fractions of the ligand with the binding pocket residues.

## 4. Discussion

It is possible to create predictive models by associating the molecular structures of ligands, either in 2D or 3D, with their biological properties, such as activity or other physicochemical characteristics. Computational biology has long relied on powerful techniques like Quantitative Structure-Activity Relationship (QSAR) models and common pharmacophore models tailored for different targets. However, these models have limitations. To develop stable, successful, and statistically significant QSAR or common pharmacophore models, a substantial amount of experimental data covering a wide range of activities related to the relevant biological property is necessary, which is not always readily available. The general concept of pharmacophore modeling stems from the fact that functional chemical groups placed in spatially and geometrically similar coordinates will induce similar responses on the same receptor, which would in turn result in similar biological activity. Particularly, feature-based pharmacophore modeling approaches match ligand substructures with common chemical features (i.e. hydrogen bond donor, hydrogen bond acceptor, aromatic ring system, hydrophobic, etc.) and positions them in a 3D special arrangement.

Static structure-based pharmacophore models carry the properties of the co-crystallized ligand in the target structure. While these models are optimized for efficiently screening small molecule libraries, they are less effective when screening libraries composed of molecules with significantly different structural features from the bound ligand. Structure-based pharmacophore modeling revolutionizes classical 3D pharmacophore hypothesis modeling by adding target information, consecutively enhancing the yield of high-throughput virtual screening studies. Latest improvements in pharmacophore modeling include the integration of 3D pharmacophore information with machine learning and artificial intelligence applications and web servers accessible to everyone. However, static structure-based pharmacophore modeling has its limitations. Notably, these models fail to account for the conformational changes and position-dependent effects of the binding cavity on ligand binding and activity. Furthermore, static models also omit the flexibility of the ligand with respect to the receptor and its conformational changes, dramatically loosing fundamental information about the receptor-ligand binding effect. This, in turn leads to an overestimation of the ligand affinity and specificity, and thus decreases the yield of pharmacophore-based virtual screening studies.

To overcome the limitations that static pharmacophore modeling may pose, we propose an approach that captures the dynamic interactions between the receptor and ligand while also utilizing the ease of application from static, feature-based pharmacophore modeling: Dynamic Receptor-based Active GeneratiON of Pharmacophores (DRAGON-P). The DRAGON-P attempts to take a step beyond simple feature-based pharmacophore modeling by taking in consideration the dynamical features of proteins and their conformational fluctuations by incorporating MD simulations to increase the sampling of the binding effects. From thereof, ensembles of structure-based pharmacophores are generated to elucidate the key ligand chemical features involved in biological activity.

The development of dynamic pharmacophore models using MD simulations offers several advantages. First, MD simulations better represent the physiological environment (e.g., pH 7.4, 310 K, 1 atm) compared to docking simulations, and they allow the tracking of pharmacophore combinations formed during the simulations. Second, these models provide a more accurate representation of the binding process by capturing the dynamic behavior of ligand-protein interactions. Third, dynamic structure-based pharmacophore models can sample multiple conformations of the ligand and protein, allowing for a more comprehensive exploration of the ligand’s binding mode. Fourth, considering the conformational changes occurring during the binding process, dynamic models provide a more accurate and reliable representation of the binding mode and can better predict ligand binding affinities. Finally, dynamic pharmacophore models can explore compound libraries with different structural properties from the co-crystallized ligand in the binding site, enabling a more extensive investigation of potential ligands.

Overall, dynamic structure-based pharmacophore models offer many advantages over static ones, providing a more accurate and comprehensive representation of ligand-protein interactions. The use of MD simulations in generating dynamic pharmacophore models allows for the inclusion of low-energy different rotamers of side chains within the binding pocket during the binding process, facilitating the creation of a more diverse sampling of ligand conformations within the binding pocket.

In this study, YB1 served as the model system, known for its multifunctional role associated with tumor progression and treatment resistance in various cancers. To target YB-1, a known inhibitor was employed, and dynamic structure-based pharmacophore models were developed for the binding pocket of YB-1. These models were subsequently applied in the virtual screening of extensive ultra-large ligand databases, resulting in the identification of new small molecule inhibitors with the potential to target YB-1 effectively.

## 5. Conclusions

YB-1, a highly conserved nucleic-acid-binding protein belonging to the CSD protein superfamily, plays a pivotal role in various cellular functions. These functions encompass transcriptional and translational regulation, DNA repair, drug resistance, and responses to extracellular signals induced by stress. Despite its critical roles, YB-1 has not received sufficient attention in research.

To address the dynamic nature of proteins and the ever-changing ligand-binding sites, we propose an alternative approach to the traditional static pharmacophore modeling. Our method involves employing dynamical pharmacophore models, which leverage MD simulations to achieve a more refined characterization of the RNA-binding site. By using dynamic, time-dependent models, we can overcome the challenges posed by the constantly shifting nature of ligand-binding sites in proteins. This refinement allows us to obtain better representative structures for virtual screening studies, offering enhanced insights into potential ligand interactions with YB-1.

