## Supplementary Figures and Tables for "Dynamic Structure-based Pharmacophore Models for Virtual Screening of Small Molecule Libraries Targeting the YB-1"

### Supplementary Information

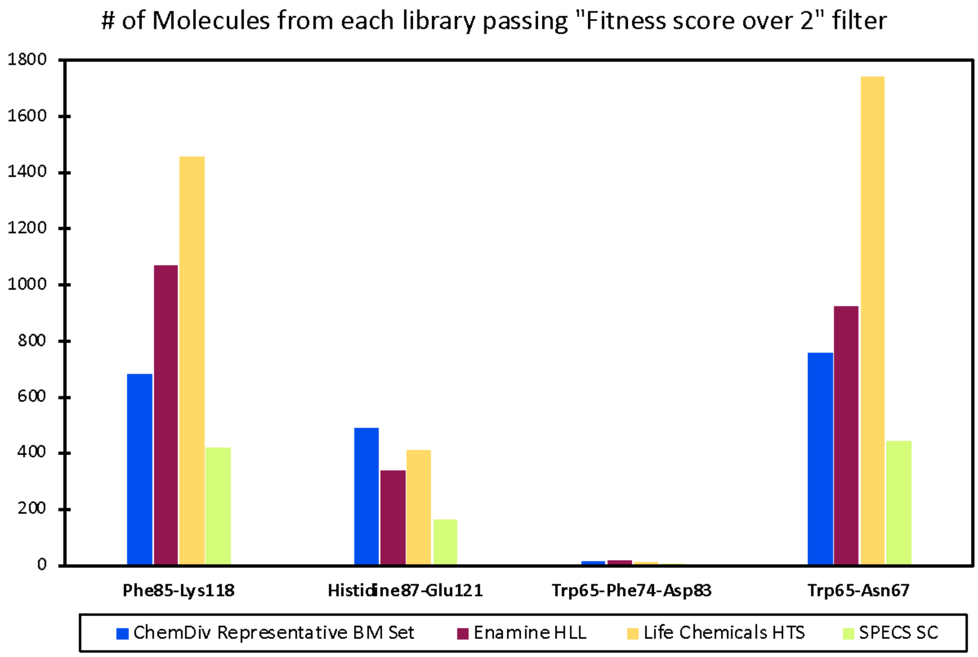


**Figure S1.** Using the optimized conformations of the protein structure and its respective ligand-receptor-derived pharmacophore hypothesis, screening of the selected four compound libraries: (i) ChemDiv, (ii) Enamine, (iii) Life Chemicals; and (iv) Specs were conducted. Only compounds that fit 3 out of 4 of the pharmacophore sites and had Phase Fitness score of over 2 were used for docking to the optimized YB-1 representative structure. The figure illustrates the number of compounds in each library that met these specified criteria.

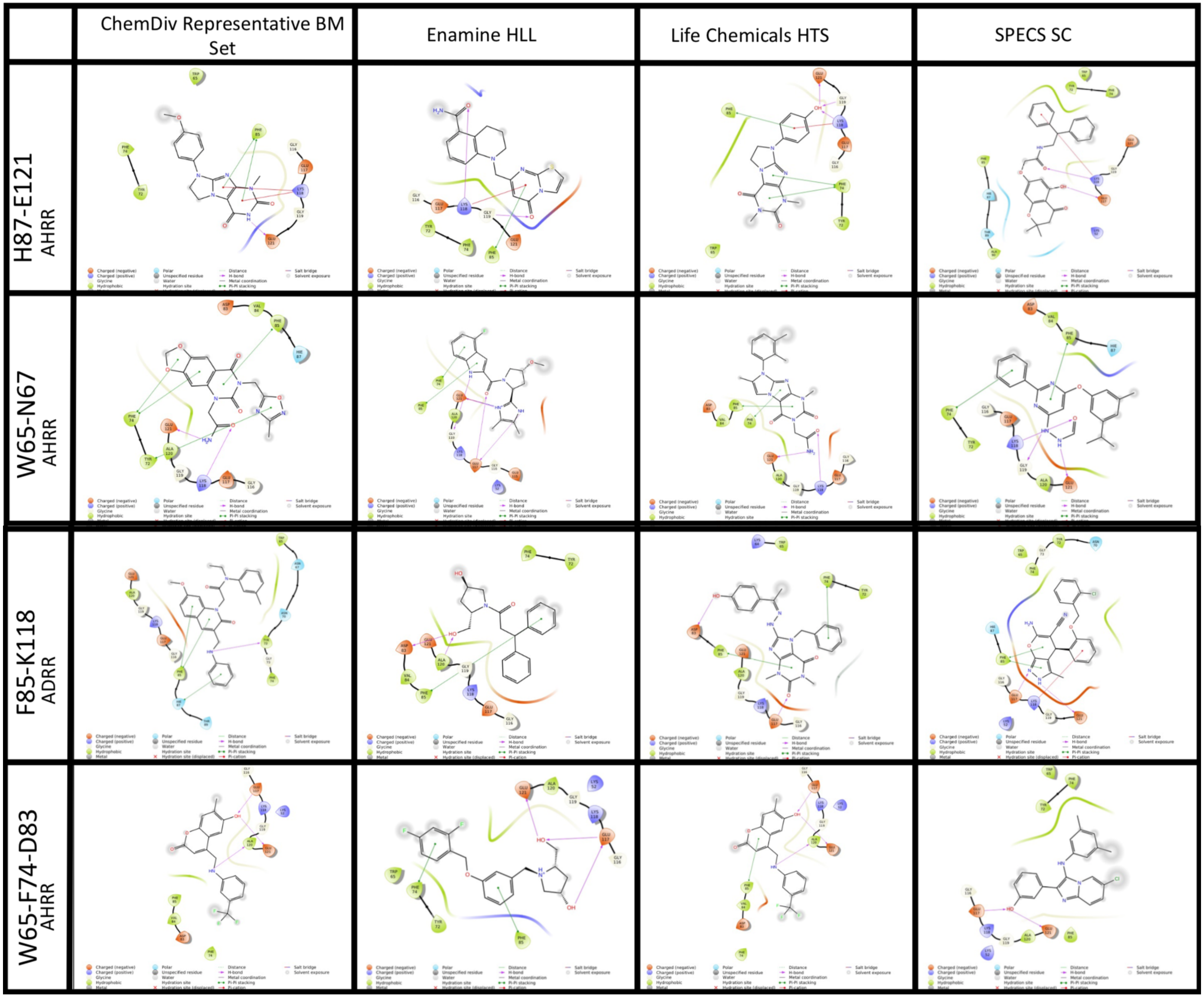


**Figure S2.** Protein-ligand interactions for selected hit compounds.
